# Astrocytic Foxo1 regulates hippocampal spinogenesis and synaptic plasticity to enhance fear memory

**DOI:** 10.1101/2023.05.01.538923

**Authors:** João Filipe Viana, Sónia Guerra-Gomes, Daniela Sofia Abreu, João Luís Machado, Sara Barsanti, Mariana Gonçalves, Cristina Martín-Monteagudo, Vanessa Morais Sardinha, Diana Sofia Marques Nascimento, Gabriela Tavares, Martin Irmler, Johannes Beckers, Michal Korostynski, Nuno Sousa, Marta Navarrete, Andreia Teixeira-Castro, Luísa Pinto, João Filipe Oliveira

**Author notes:** Corresponding author: João Filipe Pedreira de Oliveira (PhD) Life and Health Sciences Research Institute (ICVS) - School of Medicine, University of Minho, Campus de Gualtar, 4710-057 Braga, Portugal.

## Abstract

Astrocytes are active players in brain circuits, sensing and responding to neuronal activity, impacting behavior production. Activation of astrocytes triggers intracellular calcium elevations displaying complex spatiotemporal properties. Intracellular calcium activity is thought to underlie synaptic transmission, metabolism, and brain homeostasis modulation. However, the calcium-dependent signaling pathways involved in these processes are poorly understood, representing a critical knowledge gap in this field. To reveal calcium-dependent signaling pathways involved in circuit structure and function, we performed a multi-level analysis of the inositol 1,4,5-triphosphate receptor type 2 knockout (IP3R2 KO) mouse model which lacks somatic calcium elevations specifically in astrocytes. We focused on the hippocampus, a brain region responsible for cognitive function and emotional behaviors.

The transcriptomic analysis of hippocampal tissue revealed that the lack of astrocytic somatic calcium causes the differential expression of hundreds of genes. Among these, 76 genes are regulated by the astrocyte-specific Foxo1 transcription factor. This transcription factor is over-expressed in the hippocampal astrocytes of this mouse model and regulates the expression of genes involved in spinogenesis and synaptic coverage. A detailed morphological analysis of hippocampal pyramidal neurons revealed dendrites with a shift to a more immature spine profile. This spine profile shift may underlie previously described a reduction of long-term depression and performance in fear memory tasks observed in this mouse model. Indeed, we confirmed that these mice lacking astrocytic somatic calcium display an enhancement of long-term fear memory. To verify a causal relationship between these structural, synaptic, and behavioral observations, we used a viral approach to induce the over-expression of Foxo1 in hippocampal astrocytes in naïve C57BL/6J mice. This viral-driven over-expression of Foxo1 in astrocytes of the *stratum radiatum* replicated the shift to an immature spine profile in dendrites of pyramidal neurons crossing the territory of these astrocytes and led to a reduction of long-term depression in the same region. Finally, this manipulation was sufficient to enhance long-term fear memory.

The detailed characterization of the mouse model lacking astrocytic somatic calcium revealed that astrocytes modulate hippocampal circuit structure and function through Foxo1 signaling to enhance fear memory.

## INTRODUCTION

Astrocytes are active players in brain circuits, sensing and responding to neuronal activity [1,2]. This dialogue has been described to result in the modulation of neuronal activity, synchronization, and brain state shift with a consequential impact on behavior production [3–6]. Activation of astrocytes triggers intracellular calcium elevations displaying complex spatiotemporal properties [7–9]. Intracellular calcium activity is thought to underlie synaptic transmission, metabolism, and brain homeostasis modulation. The calcium-dependent release of transmitters and modulators has been extensively studied [1]. However, as with virtually every cell in the body, astrocytes display additional calcium-dependent signaling pathways, which are likely involved in the modulation of brain circuits but are still poorly understood. Among them is the calcium-dependent gene transcription to produce the machinery required and protein phosphorylation to regulate cell function and structure [10,11]. The lack of understanding of these intracellular mechanisms represents a critical knowledge gap.

To dissect calcium-dependent signaling pathways involved in circuit modulation, we studied a mouse model that lacks astrocytic somatic calcium activity. In astrocytes, somatic calcium elevations result from activating type 2 inositol 1,4,5-triphosphate receptor (IP_3_R2) at the endoplasmic reticulum. Mouse models lacking this receptor have been studied to tackle the physiological relevance of calcium signaling in astrocytes [12–27]. While astrocytes in the IP_3_R2 knockout (KO) model still display calcium activity in the distal processes and leaflets [28–30], they lack entirely spontaneous or evoked somatic calcium activity. In contrast, the neuronal activity remains intact [13,17,28], making it an excellent model for exploring calcium-dependent intracellular signaling.

This study focused on the hippocampus, a brain region deeply involved in spatial processing, cognition, and emotional behaviors. We employed complementary approaches to bridge molecular, structural, electrophysiological, and behavior analysis to reveal an astrocytic calcium-dependent signaling pathway to modulate circuit structure and plasticity with an impact on fear memory.

## RESULTS

### Lack of astrocytic somatic calcium increases Foxo1-regulated gene transcription in the hippocampus

We performed a transcriptomic analysis in hippocampal tissue from mice lacking astrocytic somatic calcium activity (IP_3_R2 KO) and respective WT littermates to dissect the astrocyte calcium-dependent intracellular pathways.

The lack of astrocyte somatic calcium activity caused the differential expression of 677 genes, of which 349 genes were significantly up-regulated (51.6%) and 328 genes were down-regulated (48.4%) (Figure 1A). This differential expression data was validated by real-time quantitative PCR (RT-qPCR) of 12 selected genes, which were chosen according to their functional relevance for cognition and astrocytic function and its fold change levels. This RT-qPCR analysis confirmed the differential expression obtained in microarray data for all up- and down-regulated selected genes (Figure 1B). To identify master regulators of the 677 differentially expressed genes, we conducted an iRegulon analysis (Table 1). This bioinformatic method analyzes the regulatory sequences associated with each differentially expressed gene to detect enriched transcription factor (TF) motifs [31]. We found nine TFs (*Foxo1*, *Vsx1*, *Sfpi1*, *Mafk*, *Stat1*, *Nkx2-1*, *Hmga1*, *Xbp1*, *Klf4*) significantly enriched based on our list of up- and down-regulated genes. Of these TFs, two – *Foxo1* and *Klf4* – display an astrocyte-enriched expression compared to other cell types, according to a published transcriptome database [32]. Specifically, *Foxo1* presented the highest enrichment score threshold (NES = 4.033) and controlled a high number (76) of differentially expressed genes.

**Figure 1.**
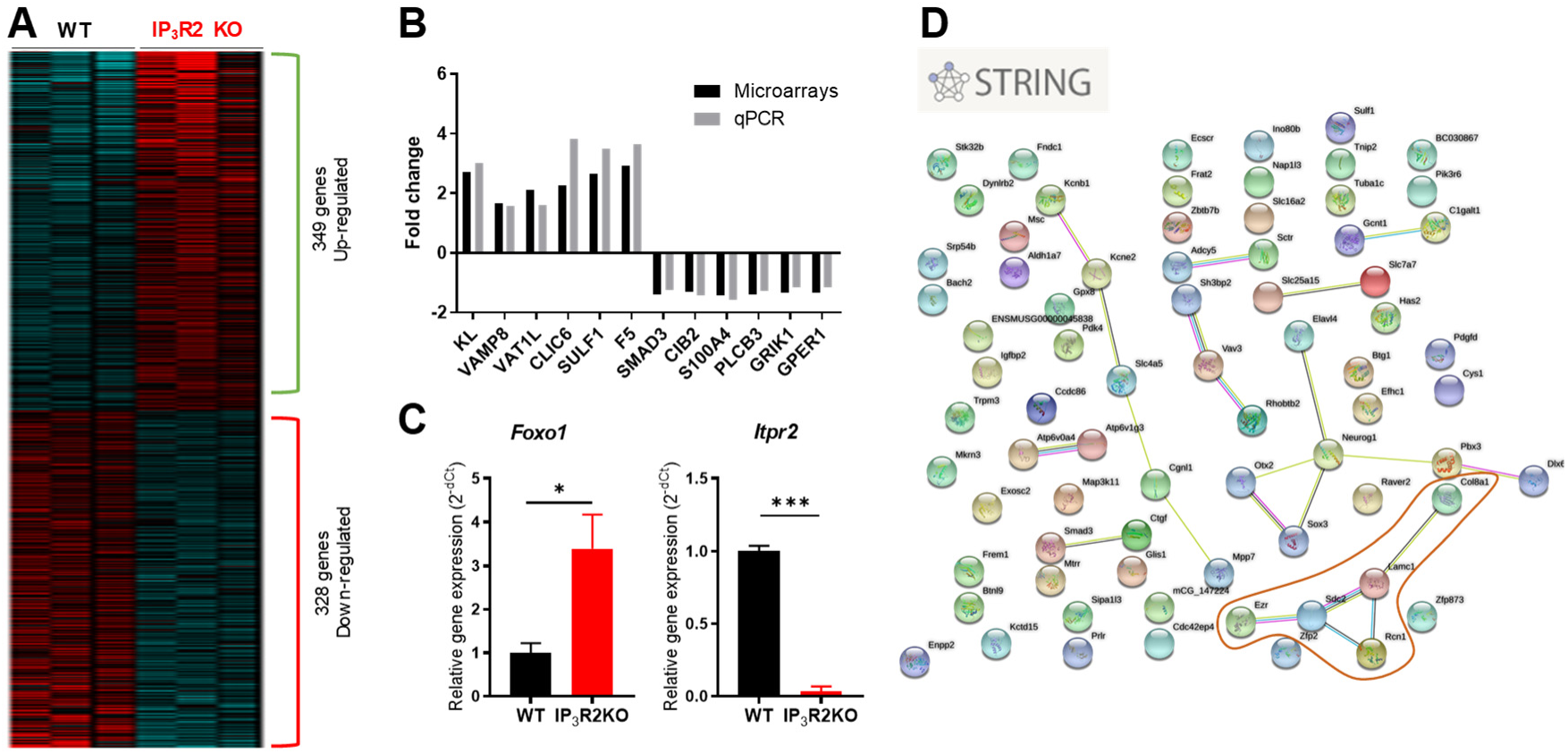
Identification of relevant molecular signaling pathways/targets related with somatic calcium signaling modulation in astrocytes. (A) Microarray analysis of the total hippocampus of WT and IP_3_R2 KO mice lacking astrocytic somatic calcium (n=3 per group) gathered 677 differentially expressed genes in IP_3_R2 KO mice as compared with WT littermates: 349 genes (51.6%) were up-regulated, 328 genes (48.4%) were down-regulated; n=3 per group). (B) RT-qPCR analysis for selected candidate genes to confirm FC levels observed in the microarray data. Black bars correspond to the observed linear FC from Affymetrix microarrays data, and grey bars correspond to the linear FC of mRNA levels calculated after normalization to 18S for each experimental group. Data presented as mean ± SEM. (C) RT-qPCR analysis for the *Itpr2* and *Foxo1* genes in an astrocytic MACS-enriched cell population obtained from the hippocampus of IP_3_R2 KO (red bars) mice and respective littermate WT (black bars). Data presented as mean ± SEM. * *p* < 0.05, *** *p* < 0.001. (D) *Foxo1* target genes (76 differentially expressed genes) were inserted in the STRING tool to look for functional interactions. This analysis revealed three clusters of genes with related functions; one cluster (in red) displays genes that are specifically enriched in astrocytes.

**Table 1.** iRegulon results regarding the most enriched transcription factors from our list of differentially expressed genes.

| TF | Name | NES | # targets | Cell type enrichment [32] |
| --- | --- | --- | --- | --- |
| **Foxo1** | Forkhead box protein O1 | **4.033** | **76** | **Astrocytes and endothelial cells** |
| Vsx1 | Visual System Homeobox 1 | 3.871 | 77 | Very low expression in the mouse brain |
| Sfpi1 | Spi-1 Proto-Oncogene | 3.573 | 19 | Microglia/macrophages |
| Mafk | MAF BZIP Transcription Factor K | 3.336 | 62 | Microglia/macrophages and endothelial cells |
| Stat1 | Signal Transducer And Activator Of Transcription 1 | 3.270 | 74 | OPCs, microglia/macrophages and endothelial cells |
| Nkx2-1 | NK2 homeobox 1 | 3.164 | 36 | Neurons |
| Hmga1 | High Mobility Group AT-Hook 1 | 3.160 | 45 | Microglia/Macrophages |
| Xbp1 | X-Box Binding Protein 1 | 3.089 | 23 | Microglia/Macrophages and endothelial cells |
| Klf4 | Kruppel Like Factor 4 | 3.043 | 10 | **Astrocytes and Microglia/Macrophages** |

Foxo1 belongs to the forkhead box O (FOXO) family of transcription factors responsible for regulating several genes involved in distinct cellular processes, with possible roles in the modulation of cognitive function [33,34]. In the brain, FOXOs are pivotal effectors of cell homeostasis, metabolism, and stress response, which may influence learning and memory [34–36]. Interestingly, the available literature reports that in different cell types [37,38], protein kinase C (PKC) controls AKT activation, which in turn is responsible for the negative regulation of Foxo1 activity [34,39]. Since PKC is tightly regulated by DAG and cytosolic calcium, in the absence of somatic calcium of IP_3_R2 KO hippocampal astrocytes, *Foxo1* activity may be disinhibited. Foxo1 is highly expressed in astrocytes in the mouse [32] and human brain [40] compared to the remaining FOXO family members. Thus, to confirm that astrocytic *Foxo1* is altered in these mice lacking somatic calcium activity, we quantified its relative expression levels through RT-qPCR in an astrocyte-enriched fraction obtained using the automated Magnetic Activated Cell Sorting (MACS). Our results show that *Foxo1* is overexpressed in astrocytes from mice that lack astrocytic somatic calcium compared to controls (Figure 1C). At the same time, the expression of the gene encoding the IP_3_R2 (*Itpr2*) is absent, as expected. We next performed a STRING analysis to explore functional associations between the proteins encoded by the 76 genes differentially regulated by Foxo1 in the hippocampus lacking calcium activity. We obtained an interaction diagram of the analyzed targets highlighting three functionally related clusters (Figure 1D). We looked for the genes in these clusters regarding their cell-type expression in the brain [32] and recognized functions described in the literature. In the microarray analysis, we found a cluster (in red, Figure 1D) including five significantly up-regulated genes (Figure 1A). Two of these genes, syndecan-2 (*Sdc2*) and ezrin (*Ezr*) are highly expressed in astrocytes [32] (Table 2). Functionally, *Sdc2* is involved in dendritic spine formation in the hippocampus [41,42], and *Ezr* teams up with *Sdc2* to link it to the actin cytoskeleton [43], with a potential role in neuritogenesis [44]. Additionally, *Ezr* is involved in filopodia formation and motility at the perisynaptic astrocyte processes (PAPs), influencing synaptic coverage [45–47]. These results suggest a regulatory role of astrocytic calcium-dependent signaling through Foxo1 of expression of genes involved in structural plasticity at the synapse.

**Table 2.** Fold change, cellular specificity, and reported functions of genes in the identified cluster.

| Gene | Fold change (microarray analysis) | Cellular specificity [32] | Described functions |
| --- | --- | --- | --- |
| **Sdc2**<br>(syndecan-2) | 1.37 | **Astrocytes and OPCs** | Dendritic spine formation [42] |
| **Ezr**<br>(ezrin) | 1.41 | **Astrocytes** | Involved in PAPs motility and neuronal filopodia formation [46] |
| Lamc1<br>(laminin subunit gamma 1) | 1.36 | Endothelial cells | Essential for basement membrane formation [62] |
| Rcn1<br>(reticulocalbin 1) | 1.38 | OPCs | calcium-binding protein in the endoplasmic reticulum (ER) [63] |
| Col8a1<br>(collagen, type VIII, alpha 1) | 2.60 | Neurons | Extracellular matrix structural component [64] |

### Mice lacking astrocytic somatic calcium activity display a shift to an immature spine profile in hippocampal pyramidal neurons

To study a putative link between astrocytic somatic calcium signaling and structural plasticity in hippocampal neurons, we conducted the tridimensional reconstruction of dorsal CA1 pyramidal neurons (Figure 2A-G). These neurons are responsible for integrating spatial, contextual, and emotional information and are actively involved in cognitive function [48]. Our results showed that CA1 pyramidal neurons of mice that lack astrocytic somatic calcium are morphologically similar to neurons of WT mice (Figure 2A-E). Specifically, the apical dendrites of pyramidal neurons of both genotypes display equivalent total dendritic length (Figure 2C) and the number of ramifications (Figure 2D). Moreover, the Sholl analysis of the dendritic tree revealed a similar intersection number at increasing distances from the soma (Figure 2E). We further analyzed the relative distribution of different spine subtypes in these neurons to assess their dependence on astrocytic somatic calcium signaling [49]. For that, we categorized the spines according to their maturity (thin, mushroom, and thick, as we previously described [50]) in proximal and distal segments of apical dendrites (Figure 2F-H). Interestingly, CA1 pyramidal neurons of mice lacking astrocytic somatic calcium display an increased percentage of thin spines (apical proximal: t_136_ = 6.69; p < 0.0001; apical distal: t_136_ = 6.36; p < 0.0001) which is accompanied by a decrease in mushroom spines (apical proximal: t_136_ = 5.40; p < 0.0001; apical distal: t_136_ = 6.15; p < 0.0001) when compared to WT neurons (Figure 2G-H). In summary, mice lacking astrocytic somatic calcium possess dorsal CA1 pyramidal neurons with normal morphology but with a shift to more immature spines. The next question is whether this structural alteration influences synaptic plasticity. Indeed, recent reports showed that mice lacking astrocyte somatic calcium signaling display decreased long-term depression (LTD) in CA1 pyramidal neurons [18,24]. Moreover, these reports link the decrease in LTD with an enhancement in the performance in the contextual fear memory task, suggesting a possible link between astrocytic somatic calcium, spinogenesis, synaptic plasticity, and fear memory processing.

**Figure 2.**
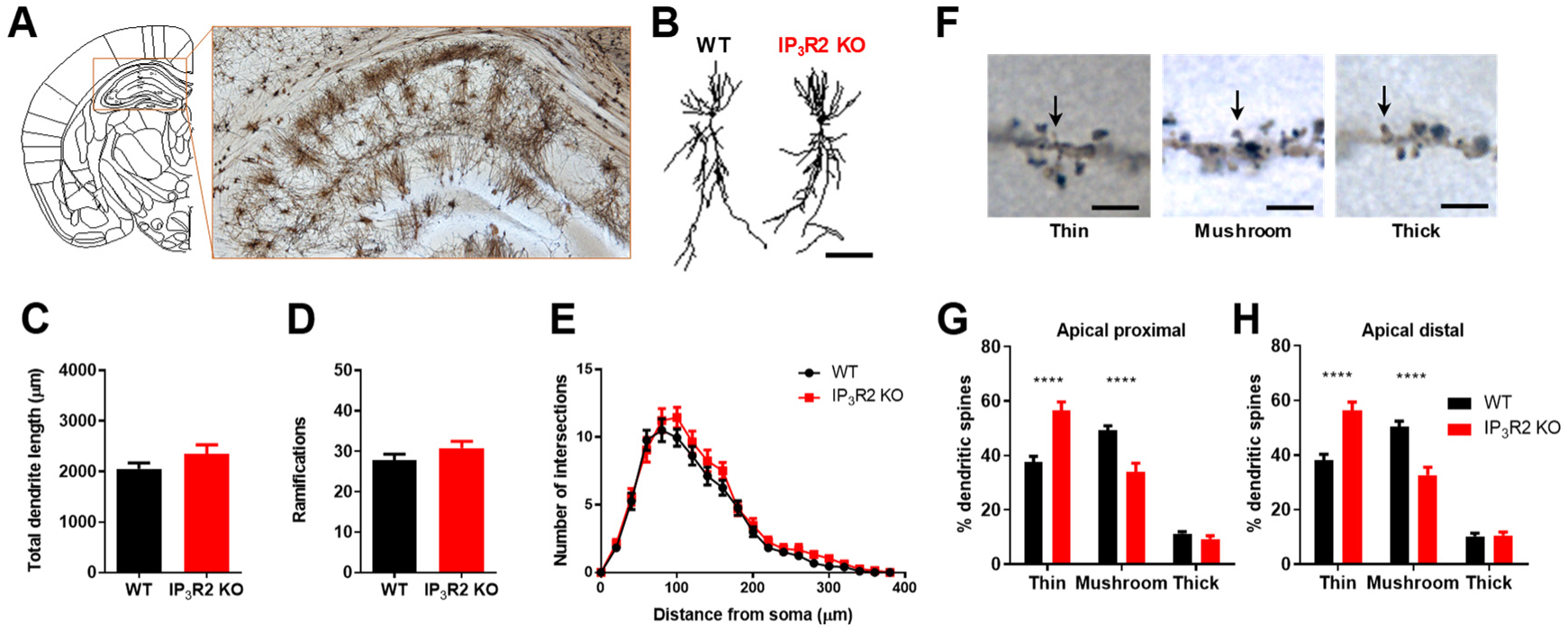
IP_3_R2 KO mice lacking astrocytic somatic calcium display more immature spines in CA1 pyramidal neurons. (A) Representative 3D reconstructions of dHIP CA1 pyramidal neurons from WT and IP_3_R2 KO mice (scale bar = 100 µm). (B-D) Morphological parameters of apical dendrites, namely total length and the number of ramifications (n=22-27 neurons per group). (E) Sholl analysis for intersections of apical dendrites. (F-H) Percentage of each spine type at apical proximal and distal dendrites; (F) Adapted from [50] n=17-19 neurons per group. WT mice are represented in black dots and lines/bars, and IP_3_R2 KO mice in red dots and lines/bars. Data plotted as mean ± SEM. * *p* < 0.05, **** *p* < 0.0001.

### Mice lacking somatic calcium signaling in astrocytes display enhanced fear memory

To explore a possible consequence of the astrocyte calcium-dependent molecular and structural changes in fear memory performance, we tested mice lacking astrocytic somatic signaling in the contextual fear memory task.

On the training day (Day 1), we found similar baseline activity and freezing responses after applying three footshocks associated with a light cue (Figure 3A). These results discarded any genotype-related alterations in baseline activity and confirmed the similar conditioning of both mice groups. Moreover, we observed that the light-shock pairings triggered a conditioned fear response in both genotypes (session, F_1,35_ = 433.2; p < 0.0001). On the following day (Day 2), re-exposure to the same context revealed a significant increase in freezing response by mice lacking astrocytic somatic calcium signaling, as compared to WT littermates (Figure 3B; t_35_ = 2.22; p = 0.03), which was not visible when mice were placed in a different context B. In the cue probe phase (Day 3), no differences between genotypes were observed after light stimulus presentation (Figure 3C), confirming that both genotypes maintained the conditioning.

**Figure 3.**
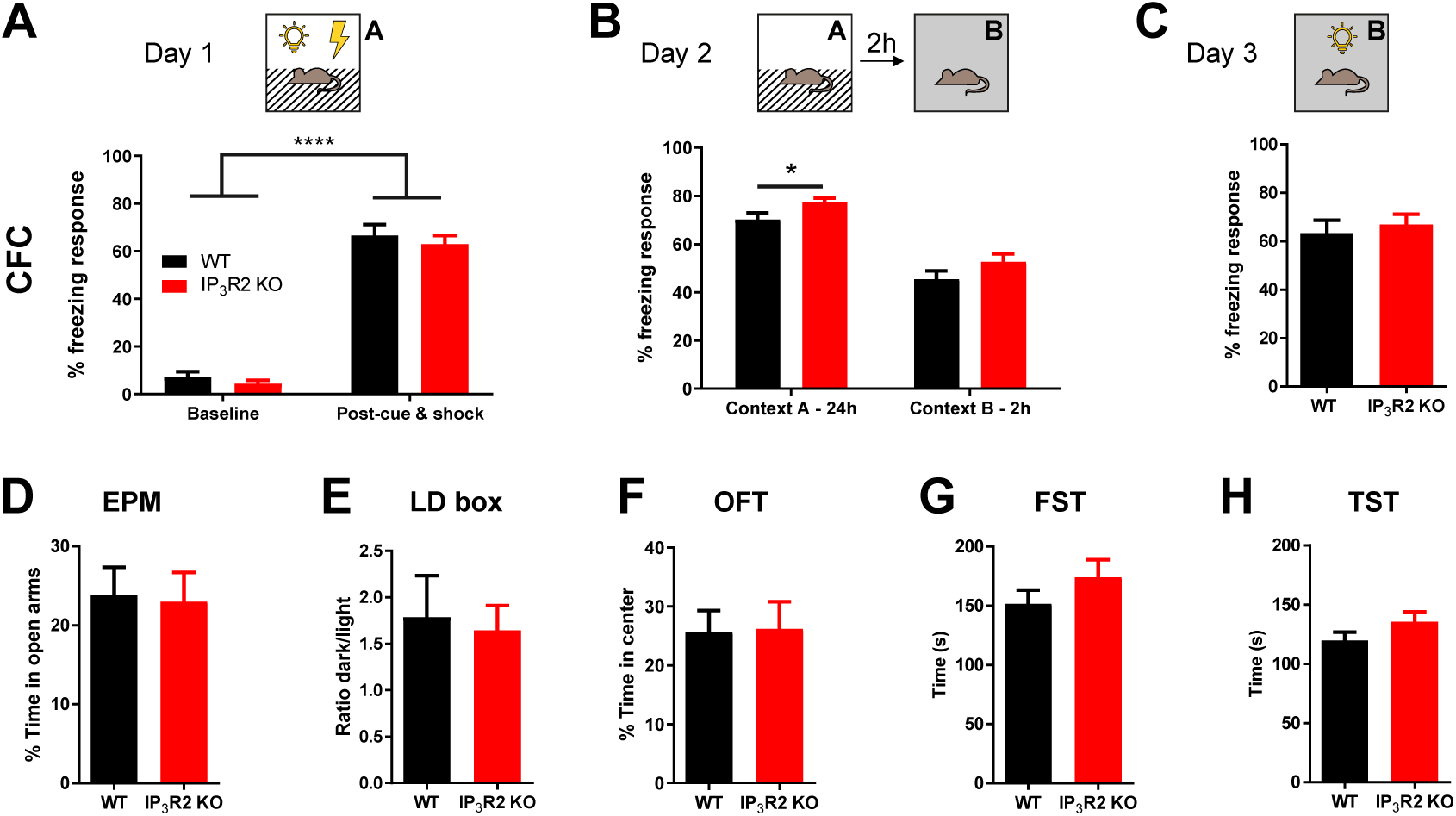
Mice lacking astrocytic somatic calcium display enhanced context-dependent fear memory and normal anxiety and depressive-like behaviors. (A-C) WT and IP_3_R2 KO mice were tested in a contextual fear conditioning paradigm (n=16-21 per group). (A) Training session depicting baseline activity (left) and freezing acquired after light-shock exposure (right) at context A. (B) Re-exposure to context A (left) enhanced freezing response in IP_3_R2 KO mice, compared to WT mice but revealed similar responses after switching to a new context (context B, right). (C) Presentation of a light stimulus in context B led to similar freezing responses between genotypes. (D) Elevated Plus Maze (EPM) reveals a similar percentage of time in open arms and the number of entries between genotypes (n=10-11 per group). (E) Light/Dark (LD) box showing an equal exploration time of dark/light of WT and IP_3_R2 KO mice (n=6-7 per group). (F) Open Field test (OFT) displays no differences between genotypes regarding time in the center (n=11 per group). (G) Forced Swim Test (FST) and (H) Tail Suspension Test (TST) allow us to evaluate latency to immobility and total immobility time for each paradigm (n=6-11 per group). WT mice are represented in black, and IP_3_R2 KO mice are in red bars. Data are presented as mean ± SEM. * *p* < 0.05, **** *p* < 0.0001.

To discard any anxious-like behaviors that could bias the behavioral performance of the mice, we performed three complementary behavioral tests widely used to assess this behavior dimension: the Elevated Plus Maze (EPM), Light/Dark (LD) box, and Open Field Test (OFT) (Figure 3D-F). Our results show that mice lacking somatic calcium in astrocytes and respective littermate controls spent a similar amount of time in the EPM open arms (Figure 3D). Moreover, both groups spent identical time in the light compartment of the LD box (Figure 3E) and the anxiogenic center of the OFT arena (Figure 3F). These results indicate that lacking astrocytic somatic calcium does not induce an anxious-like phenotype. Next, Forced Swim Test (FST) and Tail Suspension Test (TST) were performed to examine learned helplessness, indicative of depressive-like behavior, and could interfere with the behavioral evaluation. Both mice groups exhibit similar immobility times in FST (Figure 3G) and TST (Figure 3H). These observations suggest that the lack of astrocytic somatic calcium activity does not promote depressive-like behaviors.

Overall, these results support the role of astrocytic somatic calcium signaling in the fear memory performance.

### Overexpression of FOXO1 in CA1 astrocytes of the dorsal hippocampus drives fear memory enhancement in C57BL/6J mice

Until this stage, the multi-level assessment of the IP_3_R2 KO mouse model revealed that the lack of astrocytic somatic calcium signaling leads to enhanced function of the astrocyte-specific Foxo1 transcription factor. Foxo1 is over-expressed in the hippocampal astrocytes of this mouse model and regulates the expression of genes involved in spinogenesis and synaptic coverage. In accordance, we also observed a shift to immature dendritic spines in dorsal CA1 pyramidal neurons, paired with an enhancement of long-term fear memory. These observations might be linked to the reduction of long-term depression and performance in fear memory tasks observed in the *stratum radiatum* by others studying this mouse model [18,24].

Based on the above, we hypothesized that astrocytic Foxo1 could be involved in these changes to enhance the performance in the fear memory task. To test this hypothesis, we evaluated whether selective overexpression of FOXO1 in hippocampal astrocytes would influence the distribution of dendritic spines, synaptic plasticity, and fear memory in C57BL/6J mice.

To this end, we performed stereotaxic injections of an AAV5 encoding mouse FOXO1 and the mCherry reporter under the control of a GFAP(0.7) promotor. Mice were bilaterally injected in the dorsal CA1 hippocampal subregion with AAV5-GFAP(0.7)-mCherry-2A-m-FOXO1 virus (Figure 4A, Astrocyte-FOXO1+ group). As a control, a group of mice received a similar AAV5 expressing only the mCherry report gene under the control of the GFAP(0.7) promotor (Figure 4A, Control).

**Figure 4.**
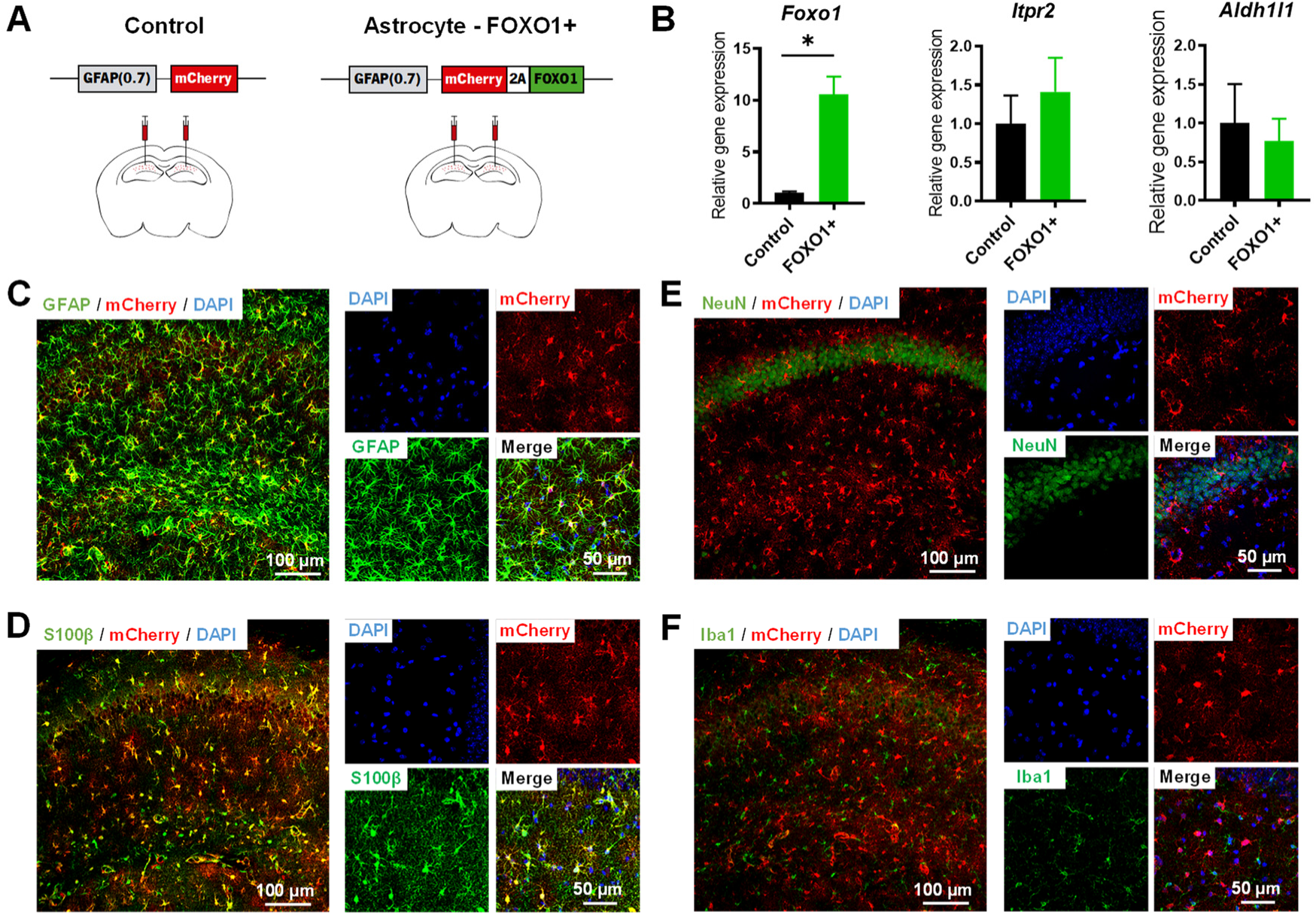
Overexpression of Foxo1 in the hippocampus is specific for astrocytes. (A) Representative scheme depicting the bilateral stereotaxic injections of AAV5-GFAP(0.7)-mCherry-2A-m-FOXO1 virus in the hippocampal CA1 area of C57BL/6J mice. (B) Quantitative RT-PCR for the Foxo1, Itp2 and Aldh1l1 genes of CA1 hippocampal astrocytes from FOXO1+ and control mice. Data are presented as mean ± SEM. * *p* < 0.05. (C-F) Confocal micrographs of immuno-histochemistry in brain slices illustrating co-expression of mCherry reporter with astrocytic (S100β and GFAP), but not neuronal (NeuN) or microglia (Iba1) markers. (C) mCherry (red) reporter gene is co-expressed with GFAP (green). (D) mCherry (red) reporter gene is co-expressed with S100β (green). (E) mCherry (red) expression is absent in NeuN^+^ neurons (green). (F) mCherry (red) expression is absent in Iba1^+^ microglia (green). DAPI staining, blue. Scale bars depicted in representative images.

We conducted a quantitative RT-PCR in hippocampal astrocytes from Astrocyte-FOXO1+ mice after autoMACS, where we confirmed a significant increase in *Foxo1* gene expression (Figure 4B). Interestingly, the expression levels of astrocyte-specific genes *Itpr2* and *Aldh1l1* were maintained in Astrocyte-FOXO1 mice (Figure 4B). To confirm the cellular specificity of the viral manipulations, we performed double-labeling of mCherry reporter in combination with astrocytic (GFAP and S100β), neuronal (NeuN), or microglia (Iba1) markers (Figure 4C-F) in brain tissue collected four weeks after the viral injection. Our results show that the mCherry reporter co-expressed with both GFAP+ (Figure 4C) and S100β+ (Figure 4D) cells, which confirms the specificity of the virus for astrocyte populations expressing these widely accepted markers. Moreover, we excluded any neuronal expression of the virus, as shown by the immunostaining for the neuronal-specific nuclear protein marker, NeuN, which never overlapped with anti-mCherry staining (Figure 4E). Additionally, the lack of co-localization of microglia marker, Iba1, and anti-mCherry staining excluded any microglial expression of the virus (Figure 4F). These results confirm that the viral transduction caused gene expression, specifically in astrocytes.

To verify a causal relationship between these structural, synaptic, and behavioral observations in the mouse lacking astrocytic somatic calcium signaling, we used a viral approach to induce the over-expression of Foxo1 in hippocampal astrocytes in naïve C57BL/6J mice, and we carried out the multi-level analysis.

To assess the involvement of astrocytic FOXO1 in the remodeling of dendritic spines in the CA1 hippocampus, we overexpressed FOXO1 in hippocampal CA1 astrocytes of Thy1-EGFP transgenic mice (Figure 5A). We found a significant increase in thin spines in apical dendrites of Astrocyte-FOXO1+ compared to mice injected with the control virus (Figure 5B). Furthermore, we observed a considerable reduction in mushroom spines (Figure 5B), resembling the previously described phenotype observed in mice lacking astrocytic somatic calcium. Spine profile shifts are generally related to changes in synaptic plasticity.

**Figure 5.**
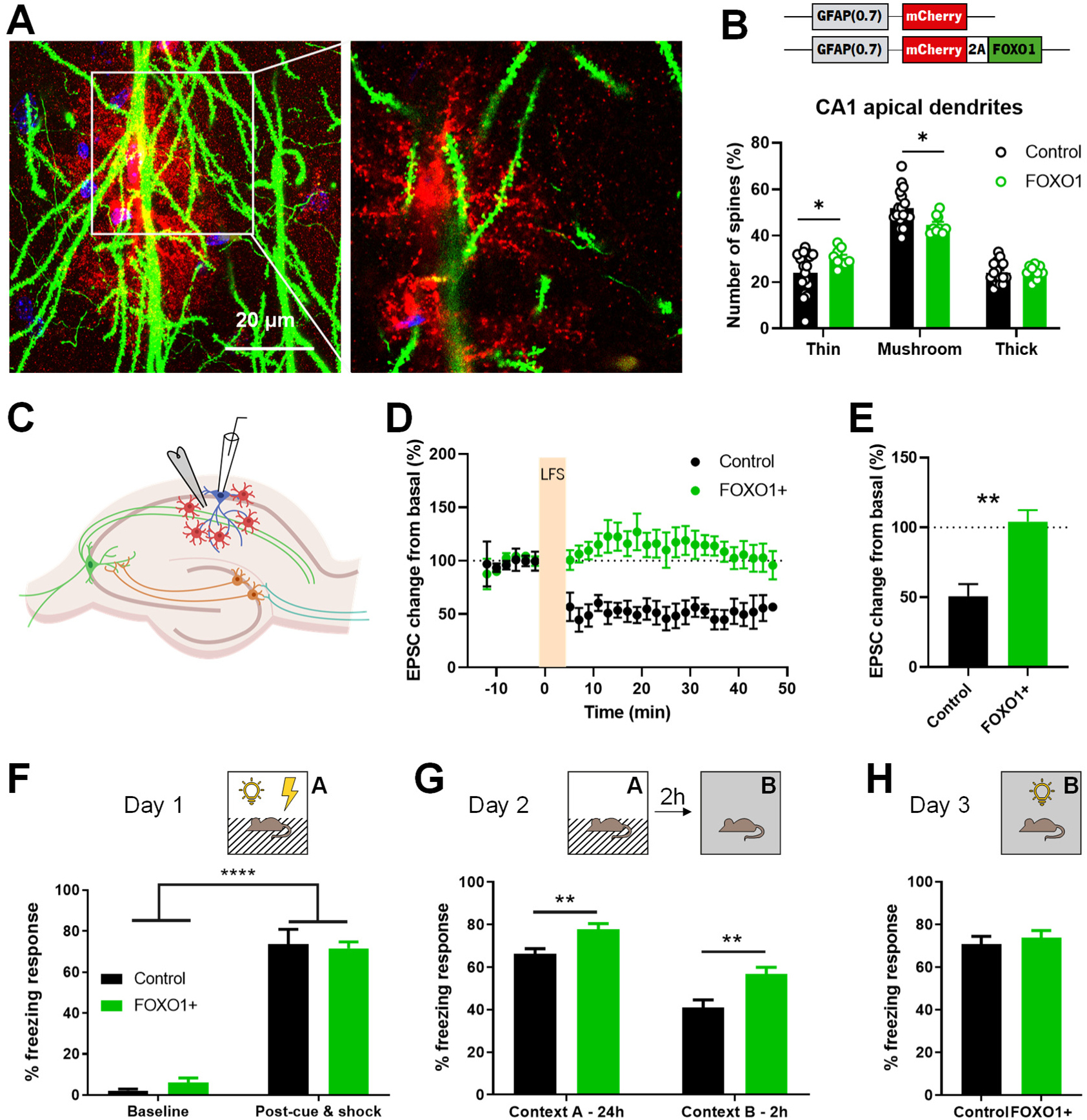
Overexpression of Foxo1 in the hippocampus is specific for astrocytes and induces alterations of dendritic spines, an impairment in LTD, and increases fear memory in C57BL/6J mice. (A) Confocal micrographs of immuno-histochemistry in brain slices illustrating an EGFP+ CA1 pyramidal neural dendrites in the territory of a mCherry+ astrocyte that overexpresses FOXO1. (B) Percentage of each spine type at apical dendrites (n=10-14 neurons per group). (C) The scheme shows the experimental configuration for electrical stimulation in *stratum radiatum* and the whole-cell recording of CA1 pyramidal neurons. (D) Relative EPSCs before and 30 min after LFS in acute hippocampal slices. (E) Average relative changes versus time during an LTD experiment with control slices (black, n=5) and slices from FOXO1+ mice (green, n=10). Time zero corresponds to the onset of the LTD protocol. (F-H) Control and FOXO1+ mice were tested in a contextual fear conditioning paradigm (n= 7-12 per group).(F) Training session depicting baseline activity (left) and freezing acquired after light-shock exposure (right) at context A. (G) Re-exposure (24 h later) to context A (left) enhanced freezing response in FOXO1+ mice. After switching to a new context, FOXO1+ mice still displayed a higher freezing response than control mice (context B, right). (H) Presentation of a light stimulus in the new context led to similar freezing responses between groups. Control mice are represented in black, and FOXO1+ mice are in green bars. Data are presented as mean ± SEM. * *p* < 0.05, ** *p* < 0.01, **** *p* < 0.0001.

As the mouse model lacking somatic calcium signaling in astrocytes displays impairment in NMDA receptor (NMDAR)-dependent long-term depression (LTD) [18,24], we performed *ex vivo* recordings of excitatory postsynaptic currents (EPSCs) from hippocampal CA1 neurons of Astrocyte-FOXO1+ and Control mice (Figure 5C). The results showed that overexpression of FOXO1 in astrocytes led to a reduction of NMDAR-dependent LTD (Figure 5D-E; t_13_ = 3.987, p = 0.0015).

Finally, we tested Astrocyte-FOXO1+ and Control mice in the Fear Conditioning Task to assess whether FOXO1 overexpression in hippocampal astrocytes would recapitulate the fear memory enhancement observed in mice lacking astrocytic somatic calcium. We evaluated contextual fear memory in both groups using the same paradigm and conditions previously described. On the first day of testing, both groups displayed a similar freezing response before and after light-shock pairings (Figure 5F). Mice from Control and Astrocyte-FOXO1+ groups showed similar freezing responses after the conditioning trials, as exposure to light-shock pairings triggered a conditioned fear response in both groups (F_1,17_ = 352.2; p < 0.0001). Interestingly, re-exposure to the same context 24 h later revealed an increased freezing response of Astrocyte-FOXO1+ mice compared to Controls (Figure 5G; t_17_ = 2.918, p = 0.0096). Moreover, the exposure of both groups to a new environment two h later decreased freezing response percentages in both groups as expected, but Astrocyte-FOXO1+ mice still presented more extended freezing responses compared to Controls (t_17_ = 3.221, p = 0.005). Nevertheless, on Day 3, the presentation of the light stimulus in the new context caused similar freezing responses in both groups confirming their similar conditioning (Figure 5H).

Altogether, these results support a causal link between FOXO1 overexpression, the shift to immature spines in hippocampal pyramidal neurons, the decrease in NMDAR-dependent LTD, and the enhancement in contextual fear memory.

## DISCUSSION

Activation of astrocytes triggers intracellular calcium elevations displaying complex spatiotemporal properties. Intracellular calcium activity is thought to underlie synaptic transmission, metabolism, and brain homeostasis modulation. However, the calcium-dependent signaling pathways involved in these processes are poorly understood, representing a critical knowledge gap in this field. To reveal calcium-dependent signaling pathways involved in circuit structure and function, we performed a multi-level analysis of the inositol 1,4,5-triphosphate receptor type 2 knockout (IP_3_R2 KO) mouse model which lacks somatic calcium elevations specifically in astrocytes. We found that the Foxo1 transcription factor is over-expressed in the hippocampal astrocytes of this mouse model and regulates the expression of genes involved in spinogenesis and synaptic coverage. A detailed morphological analysis of hippocampal pyramidal neurons revealed dendrites with a shift to a more immature spine profile, which could underlie a reduction of long-term depression and performance in fear memory tasks observed previously described for this mouse model [18,24]. Indeed, we confirmed that these mice lacking astrocytic somatic calcium display an enhancement of long-term fear memory. We next drove the overexpression of FOXO1 in hippocampal astrocytes in naïve C57BL/6J mice to verify a causal relationship between these structural, synaptic, and behavioral observations. Indeed, we confirmed that FOXO1 overexpression recapitulated the shift to immature spines, the reductions of NMDAR-dependent LTD, and the enhancement of fear memory, revealing an astrocytes-specific pathway relevant structural remodeling, LTD modulation with an ultimate impact in fear memory. This finding will be discussed considering the roles of astrocytic somatic calcium, the Foxo1 function/activity in astrocytes, and the relation between spines distribution, synaptic plasticity, and fear memory.

We took advantage of the IP_3_R2 KO mouse model since it is a widely used model of astrocyte calcium-signaling dysfunction described in the literature. These mice are reported to maintain intact calcium levels in CA1 pyramidal neurons, whereas intracellular calcium elevations in the soma and main astrocytic processes are abolished [13,17]. Astrocytes from this model still present “focal” astrocytic calcium elevations in far distant microdomains [28], possibly arising from the mitochondria [29], which are expected to play a role in the control of specific synapses or micro-circuits. A recent work descibed a viral approach to express calcium pumps under the control of a GfaABC1D promoter to extrude this ion in striatal astrocytes continuously [51]. In this study, the authors found that this approach mimics the decreased frequency of spontaneous calcium signals and reduced calcium signal amplitude in astrocyte branches observed in IP_3_R2 KO mice lacking astrocytic somatic calcium. Both models responded similarly to phenylephrine, although IP_3_R2 KO mice displayed a more considerable reduction of calcium responses. These data and the available calcium imaging observations [17] confirm that IP_3_R2 KO astrocytes cannot use reticular calcium to drive calcium-dependent pathways.

The forkhead box O (FOXO) family of transcription factors regulates several genes involved in very distinct cellular processes, including cognitive function [33,34]. Four members compose the FOXO family: FOXO1, FOXO3, FOXO4, and FOXO6. It is accepted that FOXOs are regulated via two main conserved signaling pathways: the insulin/insulin-like growth factor/protein kinase B (AKT) signaling in response to growth factors and the Jun N-terminal kinase signaling upon stress conditions. These pathways regulate FOXO transcriptional activity through post-translational modifications (PTM), namely phosphorylation. However, it is known that a higher complexity of signaling pathways and PTMs regulate FOXOs activity [33]. For instance, FOXOs can be ubiquitylated, methylated, and acetylated, affecting their subcellular location and activation state [33,52]. However, to our knowledge, the mechanisms underlying FOXO regulation in astrocytes are unknown. According to a widely used transcriptomic database [32], Foxo1 is highly expressed in astrocytes compared to the remaining family members. Interestingly, the available literature [37,38] reports that in different cell types, protein kinase C (PKC) controls AKT activation, which in turn is responsible for the negative regulation of Foxo1 activity [34,39]. Since PKC is tightly regulated by DAG and cytosolic calcium, in the absence of calcium, Foxo1 activity is disinhibited. Our results showed that in the hippocampus of mice lacking astrocytic somatic calcium, Foxo1 induces the expression of many targets. Among these targets, we found astrocyte-enriched genes such as Sdc2 and Ezr [32], which play a role in perisynaptic astrocyte processes (PAPs) morphology and dendritic spine formation [42,45,53,54] and might contribute to the observed shift to more immature spines, leading to changes in synaptic function with impact to fear memory. While our findings are observed in a model of long-term modulation of gene expression, a recent study showed that Ezr modulation is involved in astrocyte PAP retraction and fear memory consolidation [47]. Further studies should pursue this link considering the Ezr regulation by Foxo1 in a calcium-dependent manner.

Synaptic plasticity is frequently linked to changes in cognitive performance [55]. Our data shows that interfering with calcium signaling contributes to a shift to more immature (thin) spines in apical dendrites of CA1 pyramidal neurons. Thin spines are recognized as “learning spines,” whereas mushroom spines are called “memory spines” [56] mainly due to their intrinsic features. Thin spines display a more plastic and transient structure and retain biochemical signals such as calcium, which favors learning mechanisms [56]. Moreover, the glia interacts closer to thin spines associated with learning [57]. A dendritic spine simplification, from large “memory” spines to thinner “learning” spines, is associated with LTD induction [56]. Evidence points to an essential role of hippocampal LTD in memory formation and consolidation [58–61] described that astrocytes contribute to heterosynaptic long-term depression in the hippocampus. However, interfering with astrocytic calcium signaling impairs this phenomenon. The Navarrete laboratory [18] recently found that astrocytes control NMDAR-dependent LTD, supporting memory enhancement. Specifically, these authors show that the deletion of p38α MAPK from hippocampal astrocytes impairs LTD, enhancing hippocampal-associated fear memory.

Moreover, a recent work provided new insights into NMDAR-dependent LTD in IP_3_R2 KO mice with impaired calcium signaling under a C57BL/6J background [24]. This study shows that IP_3_R2 KO mice present an impaired hippocampal LTD. This observation, together with [18], supports the hypothesis that LTD impairment in mice lacking somatic calcium in astrocytes could underlie the enhancement in fear memory observed in our experiments.

In summary, the detailed characterization of the mouse model lacking astrocytic somatic calcium revealed that astrocytes modulate hippocampal circuit structure and function through Foxo1 signaling to enhance fear memory.

## ACKNOWLEDGMENTS

The authors are grateful to Prof. Alfonso Araque and Prof. Ju Chen for sharing the IP_3_R2 KO mouse line. The authors acknowledge national funding through the Foundation for Science and Technology (FCT) fellowships to JFV, JLM, DSA, SGG, VMS, and IC; IF contract to JFO (IF/00328/2015); project grants PTDC/MED-NEU/31417/2017 and POCI-01-0145-FEDER-016818; grants from Bial Foundation (037/18) and” la Caixa” Foundation (LCF/PR/HR21/52410024) to JFO; Spanish Ministry of Science and Innovation grant PID2021-122586NB-I00 to MN; FPU fellowship FPU19/01667 to CMM. ICVS Scientific Microscopy Platform, member of the national infrastructure PPBI - Portuguese Platform of Bioimaging (PPBI-POCI-01-0145- FEDER-022122, is supported by National funds through the Foundation for Science and Technology (FCT) - project UIDB/50026/2020 and UIDP/50026/2020. We thank Mareike Bamberger for excellent technical assistance.

## METHODS

### Animals

All experimental procedures were carried out in accordance with the guidelines described in Directive 2010/63/EU and were approved by the local ethical committee and the national authority for animal experimentation (DGAV 023838/2019). All mice were maintained according to the guidelines for the care and handling of laboratory animals (ad libitum access to food and water in their home cages; lights maintained on a 12 h light/dark cycle; 22 ± 1°C, 55% humidity). Male IP_3_R2 KO mice and wild-type (WT) littermates were generated as previously described [65], in a C57BL/6J background (backcrossed at least 5 times) to study the effects of the deletion of IP_3_R2-dependent astrocytic calcium signaling. These mice were supplied by Prof. Alfonso Araque (U. Minnesota, USA) [17], under agreement with Prof. Ju Chen (U.C. San Diego, USA) [12]. All mice received an ear tag at weaning age (around P21), which remained unaltered throughout the experiment and allowed us to evaluate each animal’s response blindly. The collected ear tissue was used for DNA extraction and subsequent genotyping by PCR analysis using WT (forward, ACCCTGATGAGGGAAGGTCT; reverse, ATCGATTCATAGGGCACACC) and mutant allele-specific primers (neo-specific primer: forward, AATGGGCTGACCGCTTCCTCGT; reverse, TCTGAGAGTGCCTGGCTTTT) as previously described [12]. Furthermore, C57BL/6J and transgenic Thy1-EGFP mice [Tg(Thy1-EGFP)MJrs/J, RRID: IMSR_JAX:007788] were used for the overexpression of Foxo1 in hippocampal astrocytes, as described below.

### Stereotaxic viral injections

AAV5-GFAP(0.7)-mCherry-2A-m-FOXO1 and AAV5-GFAP(0.7)-mCherry were obtained from Vector Biolabs (USA), and their titer corresponds to 3.9 x 10^12^ genome copies (GC)/mL and 3.3 x 10^12^ GC/mL, respectively. The GFAP(0.7) (gfaABC1D) promoter drives the expression of both FOXO1 and mCherry, which have a 2A linker in between for protein co-expression.

Male C57BL/6J and Thy1-EGFP mice, 8-12 weeks old, were anesthetized with a mixture of ketamine (75 mg/kg, i.p.; Imalgene 1000, Merial, United States) and medetomidine (1 mg/kg, i.p.; Dorbene Vet, Pfizer, United States), injected locally in the head with lidocaine (30 µL/0.5% lidocaine) before incision and submitted to a stereotaxic surgery for the bilateral injection of either 1 µL of an adeno-associated virus 5 (AAV5) under the control of a GFAP promoter to specifically overexpress FOXO1 in astrocytes (n=13) or an AAV5 expressing only mCherry under the control of the GFAP promoter (control group, n=8) into the dHIP (coordinates from bregma [66]: 1.8 mm anteroposterior, 1.3 mm mediolateral, and 1.3 mm dorsoventral). The injection volume and flow rate were established at 100 nL/min. At the end of the procedure, the needle was pulled up 0.1 mm, and it was left in place for approximately 5 min to allow proper viral vector diffusion. After this, mice were removed from the stereotaxic apparatus and sutured. To revert the effect of anesthesia, 1 mg/kg of atipamezole was administered. After surgery, mice received subcutaneous injections of an anti-inflammatory (Carprofeno, 5 mg/kg), an opioid for analgesia (Bupac®, 0.05 mg/kg), and vitamins (Duphalyte/Pfizer, USA). Mice were allowed to recover for four weeks before behavioral testing.

### Molecular analysis

To identify molecular signaling pathways and transcription factors altered in our mouse model of astrocytic somatic calcium dysfunction, a detailed molecular analysis was performed in RNA extracted from the hippocampus of WT, IP_3_R2 KO, Control and FOXO1+ mice (n = 2-3 biological replicates/group) that did not perform any behavioral test. A single experienced researcher conducted the hippocampus macrodissection to avoid variability.

### Isolation of total hippocampal RNA

RNA was extracted from the macrodissected hippocampi of IP_3_R2 KO mice and respective WT littermates, by mechanical homogenization, at 4°C, in 1mL of Trizol Reagent (QIAzol Lysis Reagent, Quiagen, Germantown, MD, USA) per 50-100 mg of tissue using a syringe with a 20G needle. After homogenization, the samples were left incubating for 5 min at RT. Then, per 1 mL of Trizol used, 200 μL of chloroform was added to each sample, followed by shaking for 15 s. Next, the samples were centrifuged at 12000 g for 15 min at 4°C to obtain three distinct phases: RNA aqueous phase, DNA interphase, and organic (DNA and proteins) phase. The aqueous phase was transferred to a new tube, and 500 μL of isopropanol was added and left to incubate at RT for 10 min, followed by another centrifugation step at 12000 g and 4°C for 10 min. Subsequently, the supernatant was removed, and the pellet obtained was washed with 1 mL of 75% Ethanol per 1 mL of Trizol used previously. The samples were mixed using a vortex and centrifuged at 7500 g and 4°C for 5 min. Ethanol was removed, and the pellet was left to air-dry. Finally, the RNA was dissolved in 20-50 μL of RNase-free water (Sigma) and incubated for 10 min at 60°C, followed by RNA quantification using nanodrop equipment (NanoDrop ND-1000 Spectrophotometer, ThermoFisher, USA).

### Transcriptomic analysis and bioinformatic analysis

#### Expression profiling

Total RNA (40 ng) was amplified using the Ovation Pico WTA System V2 and the Encore Biotin Module (NuGEN Technologies, Inc, San Carlos, CA, USA). Amplified cDNA was hybridized on mouse Gene 2.0 ST arrays (Affymetrix, Santa Clara, CA, USA). Staining and scanning (Scanner 3000 7G) were done according to the Affymetrix expression protocol, including minor modifications as suggested in the Encore Biotion protocol (NuGEN Technologies, Inc). Statistical analysis: Itpr2 vs. Ctrl, limma t-test; fold change (FC) > 1.3x, p < 0.05, average expression (Av) > 10, n=3 per group).

#### Prediction of transcription regulators

To identify possible regulons from our list of differentially expressed genes, the Cytoscape plugin (https://cytoscape.org/) – iRegulon – was used [31]. This plugin detects transcription factors (TFs) and their direct regulated targets based on motif discovery from a list of genes. Briefly, we imported the network file containing our gene list into Cytoscape. After selecting all nodes and edges, we performed an iRegulon analysis using the following ranking options: species and gene nomenclature (Mus musculus, MGI symbols), motif collection [10K (9713 PWMs)], no track collection, putative regulatory region (20kb centered around TSS), motif rankings database [20 kb centered around TSS (7 species)], track rankings database [20 kb centered around TSS (ChIP-seq-derived)]. The following parameters were applied regarding recovery: enrichment score threshold ≥ 3.0 and maximum false discovery rate (FDR) on motif similarity = 0.001.

#### Prediction of functional interactions

Target genes regulated by the most enriched TF from our iRegulon analysis – Foxo1 – were uploaded to the STRING tool (https://string-db.org/, version 11.0). These genes encode proteins that are involved in several cellular functions. STRING analysis retrieved possible functional associations between our targets. The minimum required interaction score established was 0.4 (medium confidence). We combined these results with a published transcriptome database (http://www.brainrnaseq.org/) [32] to identify Foxo1 targets in IP_3_R2 KO mice that have functional similarity and that are astrocyte-enriched genes.

### Analysis of enriched astrocytic fractions of hippocampus

#### Cellular suspension preparation

Astrocytes were isolated from the hippocampus of WT, IP_3_R2 KO, Control (GFAP-mCherry), and Astrocyte-FOXO1+ (GFAP-mCherry-FOXO1) mice by using the automated magnetic activated cell sorting system [67]. Each experiment was isolated by pooling the hippocampus from 3 mice. Hence, n=5 represents 15 WT, 15 IP_3_R2 KO, 15 Control, and 15 Astrocyte-FOXO1+ mice. Mice were transcardially perfused under deep anesthesia with phosphate-buffered saline (PBS). Then the hippocampus was removed, dissected, and rinsed in cold Hanks’ Balanced Salt solution without calcium chloride or magnesium chloride (HBSS[-]CaCl_2_/[-]MgCl_2_; ThermoFisher Scientific). The hippocampus was cut into small pieces and centrifuged at 300g for 2 min at 4 °C, and the supernatant was discarded carefully. According to the manufacturer’s instructions, enzymatic cell dissociation was performed using a neural tissue dissociation Kit (Miltenyi Biotec, Cologne, Germany). Then, the tissue was dissociated mechanically using a 1 mL syringe and a 20 G needle. After that, the samples were resuspended with cold Hanks’ Balanced Salt solution with calcium chloride and magnesium chloride (HBSS + CaCl_2_ + MgCl_2_; ThermoFisher Scientific, Waltham, MA, USA) and filtered through a 70 μm cell strainer (Sigma-Aldrich, St. Louis, MO, USA) to remove cell clumps followed by centrifugation at 300g for 10 min at 4 °C.

#### Myelin and debris removal

After centrifugation, cells were resuspended in PBS with 0.5% of Bovine Serum Albumin (BSA, pH 7.2) and incubated for 15 min at 4 °C with myelin removal beads II (Miltenyi Biotec, Cologne, Germany) for myelin and debris removal. After that, cells were washed and centrifuged at 300g for 10 min at 4 °C. The supernatant was removed, and the pellet was resuspended in 0.5% BSA in PBS. Then, the autoMACS® Pro Separator, using a reusable autoMACS® Column for separation, was prepared to isolate the cells automatically.

#### Microglia removal and MACS sorting of astrocytes

After myelin and debris removal, the myelin negative fraction was used to obtain the astrocytes. After centrifugation of the cell suspension at 300g for 10 min at 4 °C, the cell population was resuspended in 0.5% BSA in PBS and incubated with anti-CD11b Magnetic Microbeads, for microglia (Miltenyi Biotec, Cologne, Germany), for 15 min at 4 °C. The cells were washed, and the unbound beads were discarded after centrifugation at 300g for 10 min at 4 °C. The pellets were then resuspended and placed in the autoMACS® Pro Separator to obtain a sample without microglia. After centrifugation at 300g for 10 min at 4 °C, the supernatant was removed, and the pellet was resuspended in 0.5% BSA in PBS. Then, 10 µL of FcR Blocking Reagent was added and the cells were incubated for 10 min at 4 °C. After, cells were incubated again for 15 min at 4 °C with Anti-ACSA-2 MicroBeads, to tag astrocytes. Cells were washed with 1 mL of 0.5% BSA and centrifuged for 300×g for 5 min. The pellet was resuspended in 500 µL of 0.5% BSA in PBS and placed in the autoMACS® Pro Separator to obtain a sample enriched in astrocytes.

#### Astrocytic RNA isolation

RNA was extracted from purified astrocyte preparations using RNA-binding columns (PureLink^TM^ RNA Micro Kit, Invitrogen), followed by RNA quality assessment (2100 Bioanalyzer system, Aligent).

### cDNA synthesis and real-time quantitative PCR analysis

Total RNA from the samples used for microarray analysis (total hippocampus) and astrocytic RNA from samples obtained with MACS was reverse-transcribed into cDNA using qScript cDNA SuperMix (Quanta Biosciences, Gaithersburg, MD, USA). The primers for the selected genes of interest for microarray confirmation were designed using PRIMER-BLAST (NCBI, http://www.ncbi.nlm.nih.gov/tools/primer-blast/) (Table 3).

**Table 3.**
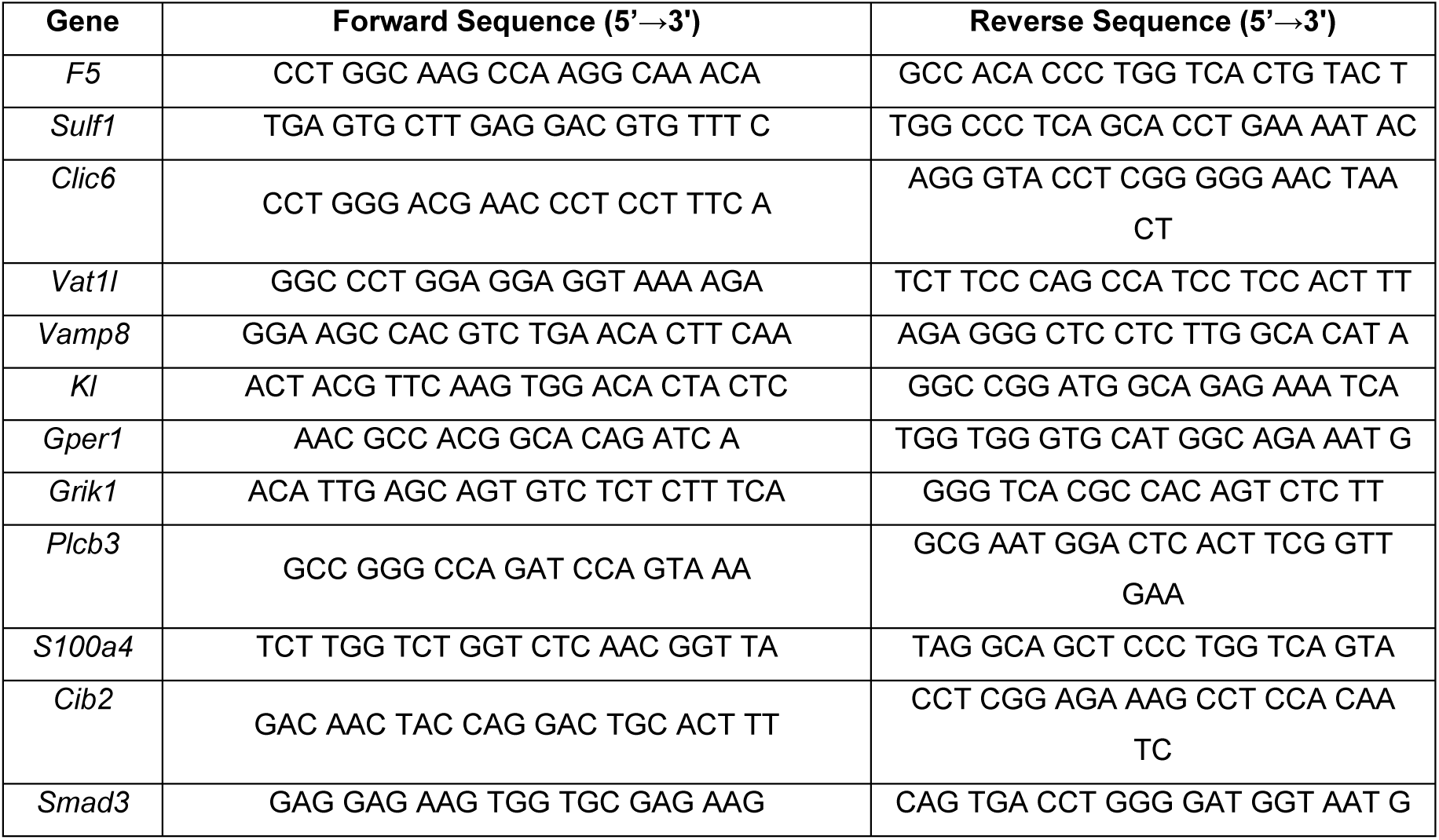
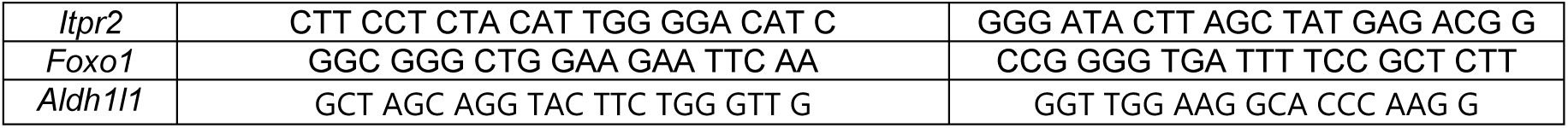
Forward and reverse sequences of oligonucleotide primers of the selected genes.

Quantifications were performed in a Fast Real-Time PCR System (Applied Biosystems, USA) using the 5x HOT FIREPol® EvaGreen® qPCR Mix Plus, ROX (Solis Biodyne, Estonia). The housekeeping genes (18S rRNA or HPRT) were used as internal controls, and the relative expression was calculated using the ΔΔCt method.

### Morphological analysis

Three-dimensional (3D) morphological analysis of neurons and dendritic spines was performed on Golgi-Cox stained neurons (for IP_3_R2 KO mice and littermate WT) and Thy1-EGFP-positive neurons (for mice overexpressing FOXO1 in astrocytes), at the CA1 region of the dorsal hippocampus. This mouse brain subregion was identified according to the mouse brain atlas [66]. Tissue processing was performed as described in the following sections.

### Golgi-Cox staining and 3D-reconstruction of CA1 hippocampal neurons of IP_3_R2 KO mice

All procedures applied to 3D-dendritic morphology were performed as previously described [50,68,69]. Mice were anesthetized with a mixture of ketamine (75 mg/kg, intraperitoneally (i.p.); Imalgene 1000, Merial, United States) and medetomidine (1 mg/kg, i.p.; Dorbene Vet, Pfizer, United States) and were transcardially perfused with 0.9% saline. Brains were impregnated in a Golgi-Cox solution (1:1 solution of 5% potassium dichromate and 5% mercuric chloride diluted 4:10 with 5% potassium chromate) for 14 days, following three washes with PBS 1x; they were transferred to a 30% sucrose solution until vibratome sectioning. Coronal sections (200 µm thick) were transferred to a 6% sucrose solution and blotted dry onto gelatin-coated microscope slides. Next, sections were alkalinized in 18.7% ammonia, developed in Dektol (Kodak, USA), fixed in Kodak Rapid Fix, dehydrated, and xylene cleared in a dark room. After this histochemical procedure, brain sections were mounted and coverslipped. CA1 hippocampal neurons from WT (n=4 mice; 22 neurons) and IP_3_R2 KO mice (n=6 mice; 27 neurons) were analyzed if they fulfilled the following criteria: 1) presence of a defined soma within the CA1 pyramidal layer; 2) complete impregnation along the entire length of the dendritic arborization; 3) no morphological alterations due to deficient impregnation or truncated branches. For each selected neuron, apical dendritic branches were reconstructed using a motorized microscope controlled by the Neurolucida software (MBF Bioscience, USA) under 100x magnification. This analysis allowed the acquisition of several parameters such as total length, number of endings, nodes, Sholl analysis, and spine number and classification. Dendritic segments of 30 µm, in the proximal and distal portions of the apical dendrite, were randomly selected to assess the percentage of each spine category (from more immature to mature: thin, mushroom, thick, and ramified) in both WT and IP_3_R2 KO mice. Data extraction was performed using the NeuroExplorer software (MBF Bioscience, USA).

### Immunohistochemistry

Four weeks post-injection, Control and Astrocyte-FOXO1+ mice (n = 2) were anesthetized with a mixture of ketamine and medetomidine and transcardially perfused with 0.9% NaCl, followed by perfusion with 4% PFA in PBS. The brain was removed and kept in 4% PFA overnight at 4°C. On the next day, it was washed in PBS 1x and cryoprotected in 30% sucrose solution. Free-floating sections (30 µm thick) were obtained using a vibrating-blade microtome Leica VT 1000 S. Sections were washed in PBS 1x and in a solution of PBS with 0.3% Triton X-100 (PBS-T, Sigma) and incubated for 1 h with a blocking solution containing 10% Normal Goat Serum in PBS-T. These steps were followed by the incubation with primary antibodies overnight at 4°C in a solution of 2% NGS in PBS 1x: chicken anti-mCherry (1:1000, HenBiotech, Portugal) and rabbit anti-NeuN (1:100, Cell Signaling, USA), rabbit anti-Iba1 (1:750, FUJIFILM Wako Chemicals, Hong Kong), rabbit anti-GFAP (1:200, DakoCytomation, USA), or rabbit anti-S100β (1:200, DakoCytomation, USA). On the next day, sections were washed in PBS 1x and incubated for 1 h at room temperature with the secondary antibodies: Alexa Fluor 594 goat anti-chicken and Alexa Fluor 488 goat anti-rabbit (1:1000, Thermo Fisher Scientific), in a solution of 2% NGS in PBS 1x. Finally, sections were rewashed in PBS 1x, incubated for 10 min with DAPI (1:1000), rinsed in PBS 1x twice, and mounted in Immu-MountTM (Thermo Scientific, USA). Images were acquired in an Olympus Fluoview FV1000 confocal microscope (Olympus, Germany) using the 20x and 100x oil immersion objectives. Moreover, Thy1-EGFP-Astrocyte-FOXO1+ mice were anesthetized with a mixture of ketamine (75 mg/kg, intraperitoneally (i.p.); Imalgene 1000, Merial, United States) and medetomidine (1 mg/kg, i.p.; Dorbene Vet, Pfizer, United States) and transcardially perfused initially with 0.9% NaCl, followed by fixation with 4% PFA in 0.1 M PBS, pH 7.4. Brains were post-fixed for 36 h in 4% PFA, and 40 μm coronal sections were cut with a vibratome. Then the slices were submitted to the same protocol described above, using the primary antibody chicken anti-mCherry (1:1000, HenBiotech, Portugal) and respective secondary antibody, Alexa Fluor 594 goat anti-chicken.

### Dendritic Spine Analysis of Thy1-EGFP-Astrocyte-FOXO1+ mice

Dendritic spines in mice overexpressing Foxo1 in astrocytes were analyzed in dorsal CA1 hippocampus pyramidal neurons using the transgenic Thy1-EGFP mice. CA1 hippocampal neurons were imaged using the confocal microscope (FV1000, Olympus, Japan). Each Z-stack image (1 µm z-step interval) was obtained with a resolution of 1024 x 1024 px under a 100x oil magnification. We randomly selected the branches of apical dendrites of pyramidal neurons (10-14 randomly selected neurons per mouse/3 mice per group), displaying GFP fluorescence throughout the entire dendritic tree enwrapped by mCherry-expressing astrocytes. The dendritic fragments and spines were traced, and the classification and quantification of spine types were performed in Fiji software (http://fiji.sc/Fiji). Each spine was then assigned to one of the classes (from more immature to mature: thin, mushroom, thick, and ramified) based on morphological criteria, as previously described [50].

### *Ex vivo* electrophysiology in brain slices

Electrophysiological recordings of CA1 pyramidal neurons were made using the whole-cell patch clamp technique in acute slices. Mice (2-3 months old, both sexes) were anesthetized by intraperitoneal injection of pentobarbital and perfused intracardially with an ice-cold solution containing (in mM) 93 NMDG, 2.5 KCl, 1.25 NaH_2_PO_4_, 30 NaHCO_3_, 20 HEPES, 25 Glucose, 2 thiourea, 5 Na-Ascorbate, 3 Na-Pyruvate, 0.5 CaCl_2_*2H_2_O, and 10 MgSO_4_*7H_2_O, bubbled with 95% O_2_-5% CO_2_. The brain was rapidly removed, and 350 μm coronal slices were made using a Leica vibratome. The slices were placed 10 minutes at 32°C in the same solution and then transferred to a recovery chamber containing acute-ACSF (in mM) 119 NaCl, 2.5 KCl, 1 NaH_2_PO_4_*2H_2_O, 11 Glucose, 26 NaHCO3, 1.2 MgCl_2_, 2.5 CaCl_2_ bubbled with 95% O_2_-5% CO2. After <1 h recovery, slices were then transferred to an immersion-recording chamber and superfused with the same gassed acute-ACSF solution. ACSF was supplemented with 50 µM picrotoxin and 5 µM CGP to block GABA_A_ and GABA_B,_ respectively. Patch electrodes were filled with the internal solution that contained (in mM): KGluconate 135, KCl 10, HEPES-K 10, MgCl_2_ 1, ATP-Na_2_ 10. In some experiments, pyramidal neurons were patched with an internal solution containing 5% biocytin. Recordings were obtained with Multiclamp 700B amplifier (Axon Instruments), digitized with Axon Digidata 1550, and pClamp software (Molecular Devices). Cells were observed under an Olympus BX50WI microscope (Olympus Optical, Tokyo, Japan) with an 40x water immersion objective. Stimulated electrodes were theta capillaries (4-9 µm tip) filled with ACSF. The stimulating electrodes were placed over SC fibers. Stimulation strength was set to elicit EPSCs of 50-100 pA in size. LTD was induced using a pairing protocol by depolarizing the postsynaptic cell to -40mV while stimulating SC fibers at 1Hz (300 pulses). The electrophysiological properties were examined prior to and at the conclusion of the experiments. The series and input resistances were monitored continuously during the experiment by applying a -1 mV pulse. The recordings were deemed stable only if the series and input resistances, as well as the duration of the stimulus artifact, did not deviate by more than 20%, to ensure reliable results. Any cells that failed to meet these requirements were excluded from the analysis.

### Behavioral testing

A behavioral assessment was performed during the light phase. One week prior to behavioral testing, mice were handled daily for 5 min. On the testing day, mice were left in the testing room for 30 min, before behavioral assessment, for habituation. Behavior tests were performed as described previously [68,70,71].

#### Contextual Fear Conditioning

Contextual Fear Conditioning (CFC) test was performed in white sound-attenuated chambers (20 cm x 16 cm x 20.5 cm) (SR-LAB, San Diego Instruments, San Diego, CA, USA) containing a light bulb on the ceiling to illuminate the entire box. The chamber where mice received a footshock (Context A) was equipped with an acrylic cylinder containing a stainless steel shock grid at the bottom. Mouse freezing behavior was recorded using a video camera and analyzed using a behavioral scoring program (Observador v0.2.7). CFC was conducted for three consecutive days and was performed according to the following paradigm:

Day 1 - Training session

Mice were placed in a white chamber containing an acrylic cylinder with a shock grid on the floor. They received three consecutive light-shock pairings of 3 min. More specifically, 20 s before the application of a foot shock, a light was turned on, and this period co-terminated with a shock of 0.5 mA. On this day, the testing room was only illuminated with a red light. This session lasted 9 min and 30 s. The first minute and the last 30 s were analyzed for the analysis of baseline and post-shock freezing response.

Day 2 - Context phase

Context A: On day 2 (24 h later), mice were re-exposed to the familiar (white) chamber where they had been conditioned, and their freezing behavior was evaluated for 3 min. A freezing response was considered when we observed complete immobility of the animal for a minimum of 1 s.

Context B: 2 h later, mice were placed in a novel context (black chamber). In this new context, room conditions changed (the room light was turned on, black plastic cardboard covered the whole box, the shock grid was removed from the acrylic cylinder, and the chamber was scented with a vanilla extract). This session lasted 3 min, and freezing behavior was scored during all session time.

Day 3 - Cue probe

On day 3, mice were re-exposed to context B conditions. After 3 min, the light bulb on the ceiling was turned on, and their freezing behavior was scored for 1 min.

#### Elevated Plus Maze

Elevated Plus Maze (EPM) test is a recognized method to assess rodent anxious-like behavior. This test consists of placing each mouse individually in the hub of a plus-like apparatus elevated 72.4 cm above the floor (ENV560; Med Associates Inc, Vermont, USA), containing two open (50.8 cm x 10.2 cm) and two closed arms (50.8 cm x 10.2 cm x 40.6 cm). Mice were allowed to explore the maze for 5 min freely. Time spent in each arm and the number of entries were recorded and analyzed using Ethovision XT 13 software (Noldus, Netherlands).

#### Light/Dark box

The Light/Dark (LD) box test is also a conventional test to assess anxiety in mice. This test is based on the innate aversion of rodents to brighter areas and their propensity to explore an arena in response to mild stressors such as a novel environment and light. The LD box apparatus consists of an arena equally divided into light and dark compartments, connected by an opening (Med Associates Inc, Vermont, USA). Mice were placed in the middle of the illuminated compartment, facing toward the dark area. Mice were allowed to explore the maze for 10 min freely. Data were extracted using the activity monitor software (Med Associates Inc, Vermont, USA), and the dark/light ratio was used as an index of anxiety-like behavior.

#### Open Field

The Open Field test (OFT) is a behavioral paradigm that evaluates motor and exploratory activity and anxious-like behavior. In this test, mice are placed in a brightly illuminated open field arena (43.2 x 43.2 x 30.5 cm) that is equipped with a system of two 16-beam infrared arrays connected to a computer. Mice are allowed to explore the arena for 5 min freely. Data were extracted using the activity monitor software (Med Associates Inc, Vermont, USA).

#### Forced Swim Test

The Forced Swim Test (FST) assesses depressive-like behavior (learned helplessness) in rodents. Briefly, mice were individually placed in transparent glass cylinders (diameter of approximately 20 cm), and filled with water at 24°C for 6 min. Each trial was recorded using a video camera and analyzed using the Ethovision XT 12 software (Noldus, Netherlands). This test is a learned helplessness paradigm based on the ability of mice to learn that there is no escape from that stressful situation. Therefore, latency to immobility and immobility time were analyzed during the last 4 min of testing.

#### Tail Suspension Test

The Tail Suspension Test (TST) evaluates mice’s depressive-like behavior (learned helplessness). This test is based on the principle that mice tend to stay immobile when they cannot escape from a stressful situation. In the TST, mice are suspended by the tail, using adhesive tape, on the edge of a shelf (approximately 80 cm above the floor) for 6 min. Each trial was recorded using a video camera and analyzed using the Ethovision XT 12 software (Noldus, Netherlands). Latency to the first immobility period and total immobility time was analyzed during the last 4 min of testing to measure learned helplessness.

### Statistical analyses

Statistical significance was considered for p < 0.05. All data assumed a Gaussian distribution as revealed by the central limit theorem for samples n > 30 (MWM data) or by the Kolmogorov-Smirnov normality test. Two-tailed Student’s t-test was applied for comparisons between WT and IP_3_R2 KO or Sham and GFAP-mCherry-FOXO1 mice. In contrast, a Two-way analysis of variance (Two-way ANOVA), followed by Sidak *post hoc* analysis, was used for multiple comparisons. The two-sided Chi-square test was carried out to analyze different behavioral categories of strategies in the MWM. All statistical analyses were carried out using SPSS 22.0 or GraphPad Prism 7.04.

Pearson coefficients were calculated for transcriptome analysis to assess the correlation between microarray and RT-PCR data. The expression console (v.1.4.1.46, Affymetrix) was used to control quality and obtain annotated normalized SST-RMA gene-level data (standard settings including median polish and sketch-quantile normalization). Statistical analyses were performed with the statistical programming environment R (R Development Core Team). Genewise differential expression testing was done using the Limma t-test and Benjamini-Hochberg multiple testing corrections (FDR < 10%). Heatmaps were generated with Genesis [72] and cluster dendrograms with the R script hclust.

